# SAUSI: an integrative assay for measuring social aversion and motivation

**DOI:** 10.1101/2024.05.13.594023

**Authors:** Jordan Grammer, Rene Valles, Alexis Bowles, Moriel Zelikowsky

## Abstract

Social aversion is a key feature of numerous mental health disorders such as Social Anxiety and Autism Spectrum Disorders. Nevertheless, the biobehavioral mechanisms underlying social aversion remain poorly understood. Progress in understanding the etiology of social aversion has been hindered by the lack of comprehensive tools to assess social aversion in model systems. Here, we created a new behavioral task – Selective Access to Unrestricted Social Interaction (SAUSI), which integrates elements of social motivation, hesitancy, decision-making, and free interaction to enable the wholistic assessment of social aversion in mice. Using this novel assay, we found that social isolation-induced social aversion in mice is largely driven by increases in social fear and social motivation. Deep learning analyses revealed a unique behavioral footprint underlying the socially aversive state produced by isolation, demonstrating the compatibility of modern computational approaches with SAUSI. Social aversion was further assessed using traditional assays – including the 3-chamber sociability assay and the resident intruder assay – which were sufficient to reveal fragments of a social aversion phenotype, including changes to either social motivation or social interaction, but which failed to provide a wholistic assessment of social aversion. Critically, these assays were not sufficient to reveal key components of social aversion, including social freezing and social hesitancy behaviors. Lastly, we demonstrated that SAUSI is generalizable, as it can be used to assess social aversion induced by non-social stressors, such as foot shock. Our findings debut a novel task for the behavioral toolbox – one which overcomes limitations of previous assays, allowing for both social choice as well as free interaction, and offers a new approach for assessing social aversion in rodents.

## INTRODUCTION

A number of mental health disorders are known to include elements of social aversion, yet, we lack the ability to adequately identify social aversion in model systems, limiting our capacity to interrogate the neurobiological mechanisms underlying this state^1,2^. While social aversion can encompass changes in social motivation, it often entails changes to a host of additional behaviors, such as social hesitancy, and social fear. Currently, there is no behavioral assay that is able to test both social motivation (social choice, interaction time, and approach) and social aversion (social freezing, reactivity, and hesitancy during free interaction), simultaneously. Various tasks have been developed to independently examine social motivation or social aversion. Tasks involving operant access to a social conspecific have been used to index social reward and motivation^3–11^. Other social motivation tasks, such as the 3-chamber sociability assay^12^, can be used to measure approach behavior and interaction time with a physically confined social conspecific^1,12,13^. In contrast, assays used to measure social aversion allow mice to freely interact with each other. These include the resident intruder assay, where a docile Balb/c intruder is placed in the home cage of the resident experimental mouse^14^. Others include free interaction between two mice in a novel arena^15^. However, none of these assays allows for comprehensive examination of both social motivation and more in depth aversion to social experience^16^.

Recent years have seen an explosion in computational tools and analysis pipelines for the estimation and analysis of complex animal behavior. For example, Social Leap Estimates Animal Poses (SLEAP)^17^, DeepLabCut^18^, Mouse Action Recognition System (MARS)^19^ are programs which utilize deep learning algorithms to track body keypoints of multiple animals. Large datasets generated by postural tracking software are then used for further supervised and/or unsupervised analysis approaches. Supervised behavioral analysis programs such as Mouse Action Recognition System (MARS)^19^ and Simple Behavioral Analysis (SimBA)^20^ allow behaviors identified by an experimenter to be automatically scored, reducing user error, bias, and differences in scoring across and even within labs^19,20^. Unsupervised behavioral analysis pipelines such as Motion Mapper^21^ and MoSeq^22^ can computationally identify patterns of motion, detecting clusters comprised of behaviors, poses, and movements that could not otherwise be detected with the human eye. More machine learning programs for behavior analysis include Stytra^23^, Anipose^24^, Janelia Automatic Animal Behavior Annotator (JAABA)^25^, DeepEthogram^26^, TRex^27^, Ctrax^28^, OptiMouse^29^, DeepPoseKit^30^, 3-Dimensional Aligned Neural Network for Computational Ethology (DANNCE)^31^, and 3D virtual mouse^32^, highlighting the speed at which innovation in this space is occurring. These tools not only reduce bias and increase efficiency, but also allow us to visualize behavioral changes in innovative and novel ways. Thus, the development of novel behavioral assays which lend themselves to these techniques is pivotal in a new age of validity, reproducibility, and translational meaning^33^.

Here, we developed SAUSI, a single, comprehensive assay which allows for assessment of motivated social choice, hesitancy, decision making, and free social interaction between pairs of mice, and which lends itself to modern machine learning and computational approaches. Because social isolation has been widely shown to reduce social motivation^13,34,35^ and other social behaviors^15,36^, we tested the impact of chronic social isolation to produce social aversion using SAUSI as our integrative behavioral readout. Using SAUSI, we found that social isolation promotes a state of social aversion, characterized by multiple factors including social fear, social hesitancy, and reduced social motivation. We demonstrated the ability to apply computational techniques to SAUSI-generated behavioral data and identify isolation state-specific behavioral motifs. We compared SAUSI to traditional assays conducted in the home cage or in a three-chamber apparatus and found that only SAUSI is uniquely able to probe multiple components of social aversion. Finally, we show that SAUSI can be used to test social aversion produced by other forms of stress, including foot shock stress, demonstrating that this assay is generalizable to various changes in an animal’s state. These experiments highlight SAUSI as a novel assay for a modern neuroscience toolkit, well equipped for the behavioral and computational analysis of complex social behaviors such as social aversion.

## RESULTS

### Selective Access to Unrestricted Social Interaction (SAUSI): a novel assay to assess social motivation and social aversion

We designed a novel arena that allows us to combine free, naturalistic interactions between an experimental mouse and a conspecific mouse (“social chamber”) and free behavior of the experimental mouse in a separate space (“home” chamber). We further designed this arena to enable assessment of motivated social behavior, such that the experimental mouse can choose to access a conspecific or withdraw from interaction via a tunnel that connects these two chambers (Fig. 1a-b). Similar to the 3-chamber task for sociability, the two chambers allow us to measure social motivation factors, such as how much time a mouse prefers to spend in the social vs. home chamber. However, instead of limiting social interaction by physically restricting the conspecific under a barrier, SAUSI allows for free interaction between the two mice when in the social chamber. Conspecifics are prevented from crossing to the opposite “home” side by receiving deterrent training prior to the social test (Fig. 1a-b). We connected the two outer chambers with a narrow tunnel, which is compatible with hierarchy testing^37^, to provide physical segregation between the two chambers, further preventing unwanted crossing by the conspecific. The tunnel allows for measurement of precise time points in the target mouse’s social decision making as they approach the social chamber, and reveals hesitancy in mice when they shelter in the tunnel for discrete periods of time or decide to back out after initiating a crossover.

**Figure 1.**
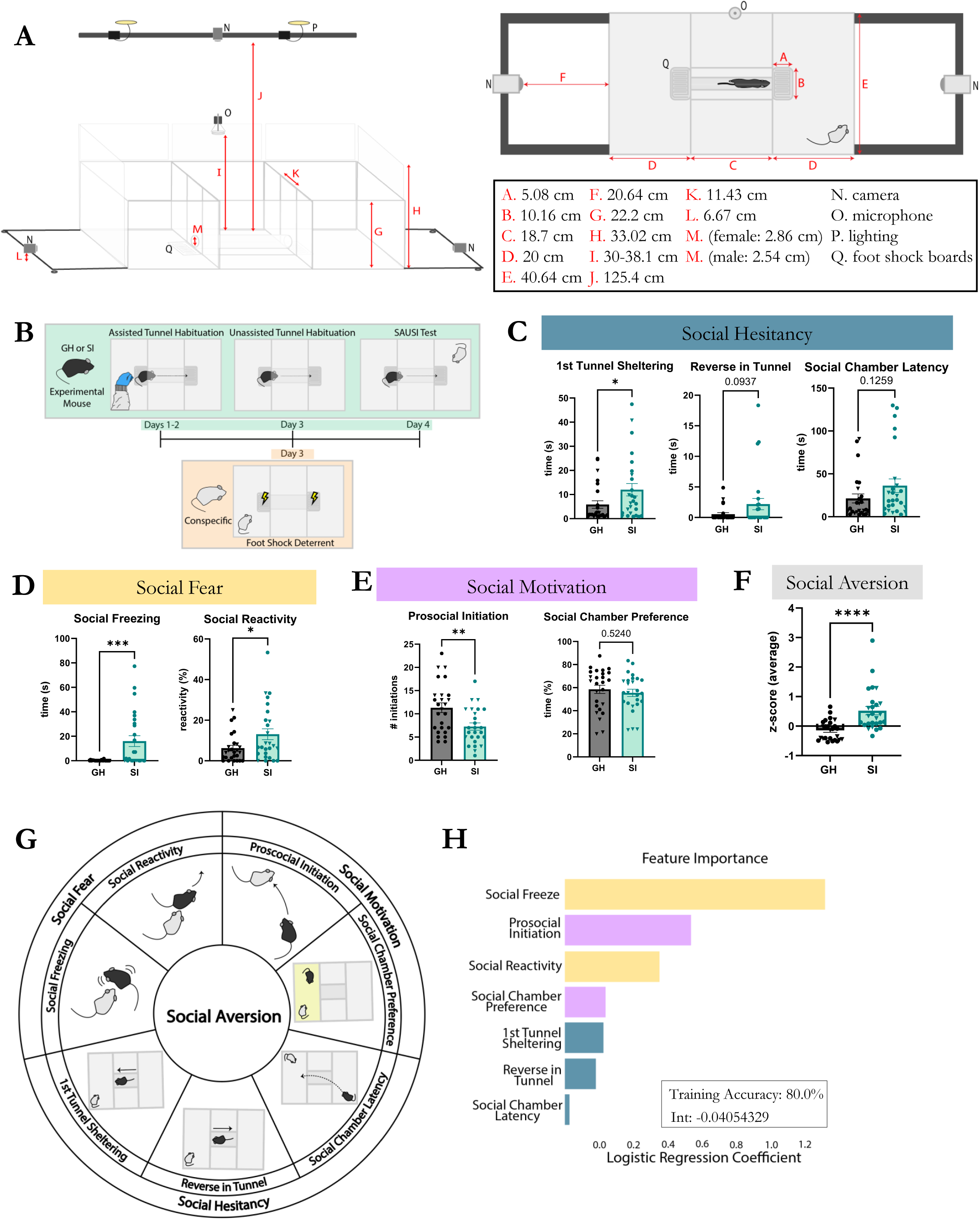
SAUSI – a novel assay to probe social aversion. (**A**) SAUSI apparatus side view (left panel) and SAUSI apparatus top view (right panel). Numbers indicate the measurements (in centimeters) for each labelled arrow in the diagrams. (**B**) SAUSI protocol. Experimental mice are habituated to the tunnel for three days (green). Conspecifics receive one day of tunnel deterrent training (yellow). (**C**) Social hesitancy was increased by isolation, with 1^st^ tunnel sheltering reaching significance. (**D**) Social fear was significantly increased by isolation. (**E**) Social motivation was negatively altered by isolation via significantly decreased prosocial initiations. No changes were found in social chamber preference. (**F**) Social aversion, represented as the z-score average of all behaviors in panel G, was significantly increased following social isolation. (**G**) Summary of SAUSI behaviors that combine to generate social aversion. (**H**) Logistic regression was performed to determine which factors are most influential to the social aversion score and the accuracy of computationally determining original housing condition based on behavior. (n=18 females (circles), n=8 males (triangles) per group; independent samples t-tests, 2-tailed).

Footshock boards were designed as the primary form of deterrent to keep conspecifics from crossing to the opposite side^38,39^. BALB/c mice were used as conspecifics, as this strain has been shown to be relatively docile, removing potential confounds related to intruder aggression^40^. To ensure that conspecific mice do not cross over into the “home” chamber or block the tunnel entryway, we employed an entry deterrent strategy, wherein a mild footshock was delivered via an electric board directly in front of the tunnel (Fig. 1a). Footshock boards were connected to grid scramblers (Med Associates), with mice completing the loop if at least one limb touched the positive and one limb touched the negative components of the footshock board (Figure 1 – figure supplement 1). To determine the bare minimum shock intensity (mA) required to serve as a deterrent without significantly altering the conspecifics’ behavior, we conducted a behavior response curve experiment (Figure 1 – figure supplement 2a). BALB/c mice were shocked at either .1mA., .2mA, .3mA or .5mA whenever they touched the footshock board across a 5-minute training session. Mice were given 1 session/day for 2 days and then tested with a group housed C57BL/6 mouse on the third day to assess social behavior and efficacy of the foot shock training (Figure 1 – figure supplement 2a). We found that .5 and .3 mA shocks, but not .1mA or .2mA, were sufficient to act as deterrents by significantly reducing the amount of time and latency of stepping on the foot shock board (Figure 1 – figure supplement 2b). However, .5mA resulted in significant changes to conspecific behavior, such as increased freezing (Figure 1 – figure supplement 2c). In contrast, .3mA worked as a deterrent while not significantly impacting conspecific behavior (Figure 1 – figure supplement 2c). There were no changes in sniffing, thigmotaxis, or aggression across all groups. Locomotion was reduced by all levels of shock, even those that weren’t effective as a deterrent (Figure 1 – figure supplement 2c). Critically, .3mA shock resulted in no changes to behavior of the experimental mouse when interacting with the mildly shocked conspecific Figure 1 – figure supplement 2d).

Many neuroscience experiments employ tethering systems to allow for neural manipulations or recordings during behavior. Because of this, we also adapted SAUSI to accommodate head mounts and tethering by developing an open-top tunnel design. This features a U-shaped channel in between the two outer chambers instead of an enclosed tunnel. This has been shown to work with wired miniature microscopes (Inscopix) as well as tethering for optogenetics experiments (Supplemental Video 1).

Collectively, this design allows for the assessment of a) free social interaction, b) solitary behaviors, c) social motivation, d) social hesitancy and social decision making, and e) general fear (baseline thigmotaxis and freezing). All of these features combined enable a robust paradigm for assessing social aversion wholistically.

### Prolonged social isolation promotes social aversion, as revealed by SAUSI

Social isolation (SI) is known to promote social aversion^13,15,34,36,41^. Yet, this has been difficult to test in rodent models due to the lack of an appropriate, wholistic assay that can measure multiple features of social aversion^1^. Thus, using SAUSI, we tested whether prolonged SI results in increased social aversion in mice. We found that isolation significantly increased social hesitancy behaviors such as time spent sheltering in the tunnel upon the first approach (Fig. 1c). Another hesitancy score, reversing out of the tunnel once the decision was made to cross over to the social chamber, trended towards significance (p=.09) (Fig. 1c). We also found that prolonged social isolation produced an increase in social fear behaviors^1^ such as social freezing (freezing in response to being sniffed by the conspecific) and social reactivity (jumping or darting in response to sniffing) (Fig. 1d)^41–43^. Additionally, social isolation caused a reduction in prosocial initiation (Fig. 1e) while preference for the social chamber remained unaltered (Fig. 1e). Finally, isolation increased the tonality of ultrasonic vocalizations (USVs), but changes in slope and total number did not reach significance (Figure 1 – figure supplement 3a). Aggression was significantly increased following social isolation (Figure 1 – figure supplement 3b), while no changes were found with sniffing, generalized anxiety measures, or mobility (Figure 1 – figure supplement 3c-e). Collectively, these results reveal multiple social aversion behaviors that are induced by social isolation and identified using SAUSI.

Next, we sought to visualize the behavioral changes observed as a wholistic, cumulative social aversion score. We converted the datasets for each behavior in the categories of social fear, social hesitancy, and social motivation into z-scores and calculated the average z-score as an index of social aversion (Fig. 1f). This social aversion z-score was significantly higher in socially isolated mice compared to group-housed (GH) controls (Fig. 1f-g). We then performed multi-logistic regression analyses and were able to determine whether a mouse was group housed or isolated with 80% training accuracy. The coefficients of this logistic regression equation reveal the behavioral features that most heavily mediate our social aversion score (Fig. 1h). The components which influenced the social aversion score most strongly were social freezing and reduced prosocial initiation, indicating that social fear and social motivation behaviors are the most important factors contributing to social aversion in isolated mice (Fig. 1h).

### Computational approaches + SAUSI can detect the social state of an animal

Social behavior in rodents is highly complex and dynamic, continually shifting as animals engage in back-and-forth interactions. Recent advances in machine learning and computational approaches for the analysis of social behavior have enabled a deeper, broader, and less biased approach towards understanding social behavior^33,44–48^. To test whether SAUSI lends itself to modern computational approaches for postural tracking and analysis of behavior, we applied a variety of computational tools to videos obtained from our SAUSI assay (behavior displayed in Fig. 1).

Thirty-six high speed videos were loaded into Social LEAP Estimates Animal Poses (SLEAP), which allows for the identification and pose estimation of multiple behaving mice^17^. We tracked 8 body points on each mouse using SLEAP (Fig. 2a) and cleaned the data to include only 2 tracks. After initial tracking was performed, blind experimenters manually corrected any switched tracks, and interpolated the videos. Next, we extracted pose estimation data and tested whether computationally extracted features could distinguish the “social state” of the animal (SI vs. GH conditions). To do so, we used Motion Mapper^21^, which enables the visualization of changes in feature space identified by stereotyped patterns of motion (Fig. 2a). Using the Motion Mapper pipeline, x/y coordinates for each body point extracted from SLEAP were processed using a series of dimensionality reductions. The raw data was converted into angles, normalized, and analyzed using principal component analyses (PCA) for the first dimensionality reduction. Next, we used UMAP to compute distances in high-dimensional space and neighboring points and embedded them closely on a new low-dimensional space, allowing for visualization of patterns of motion. Finally, we used watershed segmentation to separate the continuous stereotyped movements into groups (or regions), which were identified as collections of features (Fig. 2a). Motion Mapper identified 10 distinct regions using data from all groups combined (Fig. 2b). Occupation of feature space was visualized for each group (Fig. 2c) and time spent engaging in specific regions was quantified, revealing increased time in regions 9 and 10 for isolated mice (Fig. 2d). Next, to test whether housing condition could be decoded based on motion-mapper generated features, we employed multi-logistic regression analyses and determined housing condition accuracy and importance of each feature (Fig. 2e). Using the multi-logistic regression equation, we were able to determine whether a mouse was group housed or socially isolated with 75% training accuracy. The coefficients of this equation revealed regions 10 and 9 to be the most impactful in distinguishing between GH and SI mice. This demonstrates that the state of the mouse (GH or SI) can be mathematically predicted from unbiased, unsupervised feature analysis based on postural tracking data. Finally, using SLEAP tracking, we assessed distance between the nose of the experimental and conspecific mice and found a numerical increase for isolated mice over the course of the test phase (Fig. 2f). When assessing how behaviors change in congruence with nose-nose distance, we found a stark contrast between isolated mice and controls. Isolated mice display more social hesitancy behaviors prior to engaging in social interaction (latency to approach, longer time sheltering in tunnel on the first approach, reversing in the tunnel) and as mice gain proximity, the behaviors switch to social fear – highlighted by the presence of social freezing behaviors (Fig. 2g). In sum, the unbiased identification of stereotyped motion by unsupervised machine learning approaches revealed distinguishable GH and SI internal states based body keypoints tracked during SAUSI. Combining this with supervised behavioral analysis techniques revealed that these differences are likely due to alterations in social aversion following social isolation.

**Figure 2.**
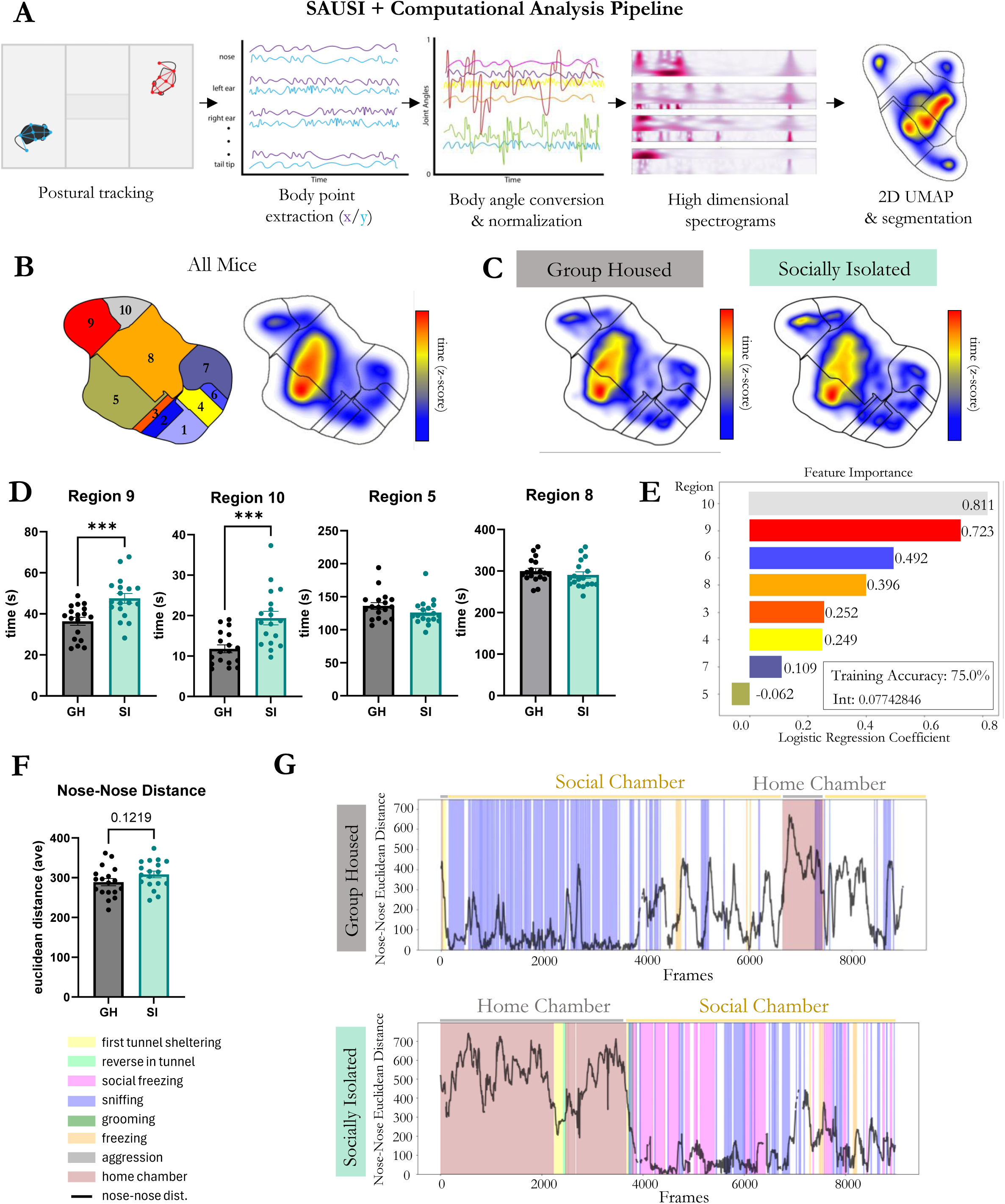
Unique behavioral footprints are detected using deep learning. (**A**) The computational workflow for unsupervised behavioral classification. (**B**) Map of behaviors that were generated by the Motion Mapper pipeline. (**C**) Heatmap distribution of time on the behavior map for GH and SI mice. (**D**) Time spent in Regions 9 and 10 were significantly increased in SI mice compared to GH controls. Other regions of behaviors had no differences between the groups. (**D**) Multi-logistic regression was performed on all behaviors identified by the Motion Mapper pipeline. Feature importance is shown with logistic regression coefficients. (**E**) Nose to nose distance was not significantly different between GH and SI mice. (**F**) Raster plots from a single animal in each group, depicting manually scored behavior and nose-nose distance during SAUSI. (n=18 females per group; independent samples t-tests, 2-tailed).

### SAUSI uniquely reveals social aversion when compared to traditional assays

To test whether SAUSI is unique in its ability to allow for the assessment of social aversion or whether social aversion could be comprehensively probed using other, well-established behavior assays, we compared behavior on SAUSI to both behavior in the 3-chamber social interaction assay^49,50^ and the Resident Intruder Assay^14,51^. In the 3-chamber social interaction assay^49,50^, experimental mice were placed in the center chamber of a 3-chamber arena. A sex- and age-matched conspecific was placed in one chamber under an overturned cup, while an inanimate object (e.g. plastic block) was placed under a cup in the opposite chamber. Given limited social interaction due to the restrictive nature of the 3-chamber design, one is unable to assess fear specific behaviors during this assay. However, we were able to see a reduction in social interaction preference and social approach preference (Fig. 3a). There was no significant difference in social interaction latency in the 3-chamber assay (Fig. 3a), similar to what was observed in SAUSI (Figure 1 – figure supplement 3a). In addition, mice vocalized in the 3-chamber (Fig. 3a), but with far fewer USVs observed than in SAUSI (Figure 1 – figure supplement 3c). There were no significant differences in call slope or other call features, in contrast to effects observed using SAUSI (Figure 3 – figure supplement 1a). This data reveals the importance of the free interaction featured in SAUSI, in that while we were able to measure one aspect of social hesitancy and social motivation, all other social fear-related behaviors were largely absent. Furthermore, limiting social interaction with physical restriction altered the number and quality of USVs in mice. These differences in social communication could impact the behavioral dynamics between two mice, further distancing them from a naturalistic social setting^52–55^.

**Figure 3.**
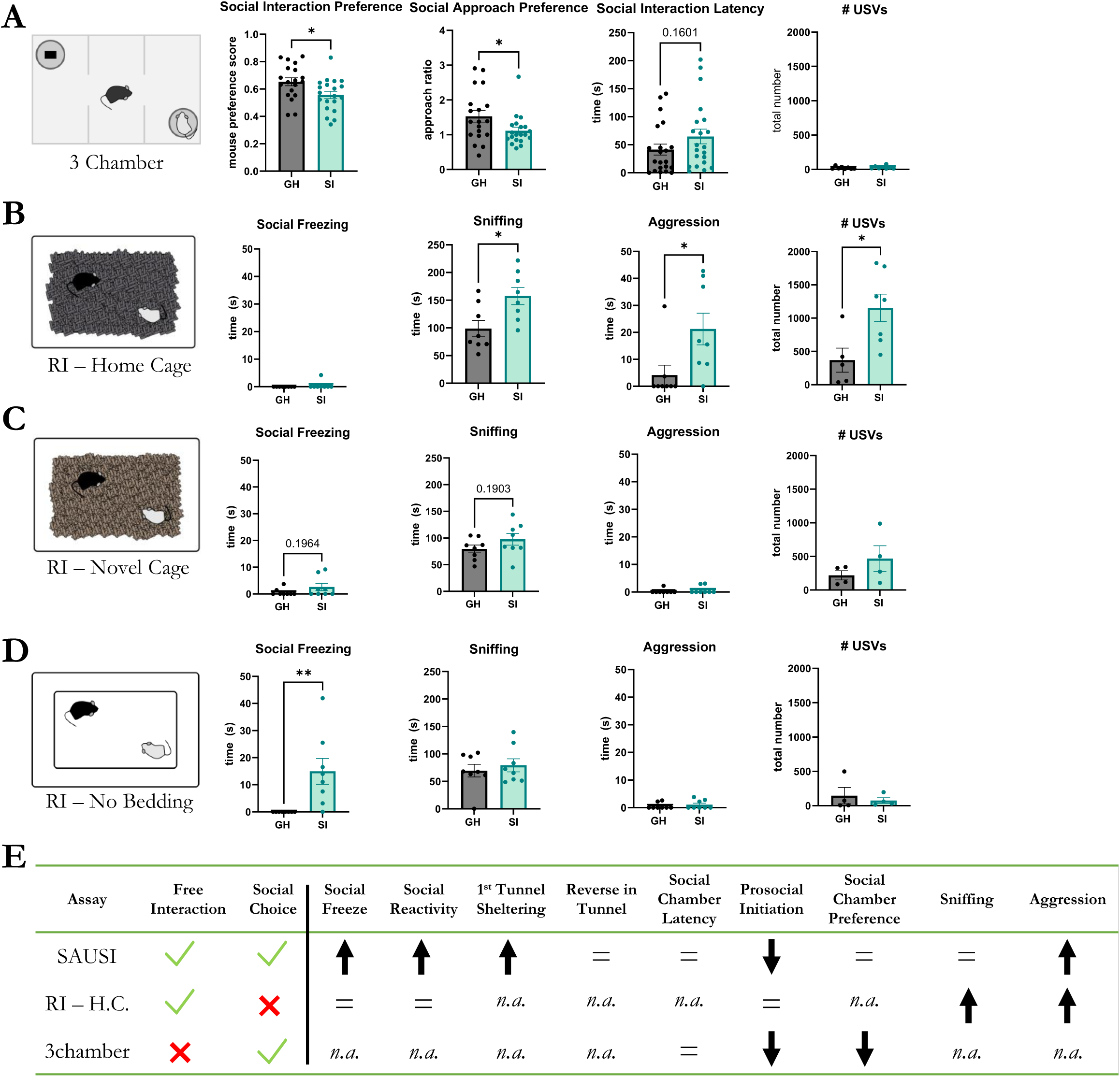
Traditional assays fail to detect comprehensive social aversion behaviors. (**A**) During the three-chamber test, isolated mice showed a reduction in social interaction preference 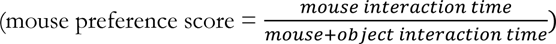 and social approach preference 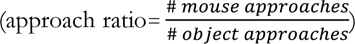, but no significant changes in latency to approach the mouse cup. Vocalizations were fewer than in other tasks but are unaltered by isolation. (n=21 females per group; independent samples t-tests, 2-tailed). (**B**) During home cage resident intruder (RI), isolated mice did not display social freezing, but did engage in higher amounts of sniffing, aggression, and vocalizing. (n=8 females per group; independent samples t-tests, 2-tailed) (**C**) During RI-novel cage, no significant differences in behavior (n=8 females per group; independent samples t-tests, 2-tailed) or USVs (n=4 females per group; independent samples t-tests, 2-tailed) were found. However, small amounts of social fear behaviors were present in isolated mice. (**D**) During RI-no bedding, isolated mice displayed social freezing behavior, but no increases in aggression (n=8 females per group; independent samples t-tests, 2-tailed). No differences were found in number of USVs made (n=4 females per group; independent samples t-tests, 2-tailed). (**E**) Summary table of the advantages of SAUSI over other assays and how behaviors differ across each test. *n.a.* means *not applicable*.

Next, we examined whether we could identify isolation-induced social aversion in a free social interaction assay, the Resident Intruder (RI) assay, another widely-used behavior assay which takes place in the home cage of the experimental mouse^51,56^. Experimental mice (GH vs. SI) were each tested with a 3-minute baseline (no intruder) and 10-minute test phase (+ Balb/c intruder). In contrast to results obtained using SAUSI, we found a lack of social fear behaviors (Fig. 3b and Figure 3 – figure supplement 1b). We also found no changes in prosocial initiation (Figure 3 – figure supplement 1b). Instead, we found a significant increase in sniffing and aggression following SI (Fig. 3b). There was also a significant increase in the number of USVs produced by isolated mice (Fig. 3b), however changes to slope or tonality were not significant (Figure 3 – figure supplement 1b). We hypothesized that the failure to find any SI-induced changes in social fear behaviors may be due to competition with territorial behaviors (sniffing and aggression)^57^, as well as the inescapability of the environment^58^.

To determine whether territorial, antisocial behavior competes with aversion behavior to eliminate social fear in the RI assay, we next performed RI in a novel cage (with new bedding), as RI-aggression has been shown to be highly context-dependent^56,58^. We found that in a novel cage environment, isolation did not elicit distinguishable features that were observed in other assays (Fig. 3c,e & Figure 3 – figure supplement 1). Indeed, while social freezing and aggression were minutely present in SI mice during novel cage testing (Fig. 3c), the magnitude was smaller than observed in other assays (Fig. 1c, Figure 1 – figure supplement 3, Fig. 3b). To test whether this could be due to the presence of bedding material, as opposed to plastic flooring (used in SAUSI), we tested animals on the RI assay in a novel cage without bedding. RI-novel-no bedding revealed significant increases in social freezing behavior (Fig. 3d), but no other significant differences were found (Fig. 3d and Figure 3 – figure supplement 1). This indicates that the absence of bedding may be a driving force of social freezing.

### SAUSI reveals social aversion across different forms of stress

To assess whether social aversion is detectable with SAUSI amongst an alternative form of non-social stress, we tested our novel assay with male and female mice that experienced foot shock stress (n=16) (vs. no-shock controls (n=16)) (Fig. 4a). Consistent with stress effects more broadly, mice that experienced foot shock stress spent significantly more time freezing during the foot shock stress paradigm than no-shock controls^59^ (Fig. 4b). When tested for social aversion using SAUSI, we found that shocked mice show significantly increased social aversion (Fig. 4c). Further analyses of social aversion sub-categories revealed that social hesitancy was significantly increased in shocked mice, specifically in the latency to enter the social chamber and time spent reversing in the tunnel (Fig. 4d). Surprisingly, and in contrast to social isolation, shocked mice did not display social fear behaviors, with almost none of the mice showing social freezing, and no differences in social reactivity (Fig. 4e). Finally, social motivation was significantly reduced in shocked mice, as shown by reduced percentage of time spent in the social chamber and in reduced number of prosocial initiations (Fig. 4f). While a decrease in baseline mobility was found in mice that received shock, there were no changes to USVs, aggression, sniffing, or generalized anxiety (Figure 4 – figure supplement 1a-e). Ultimately, SAUSI is sensitive to social aversion across different forms of stress. This sensitivity also reveals nuanced differences in aversion and motivation behaviors that differ across different forms of stress, highlighting the importance of the nature of stressor.

**Figure 4.**
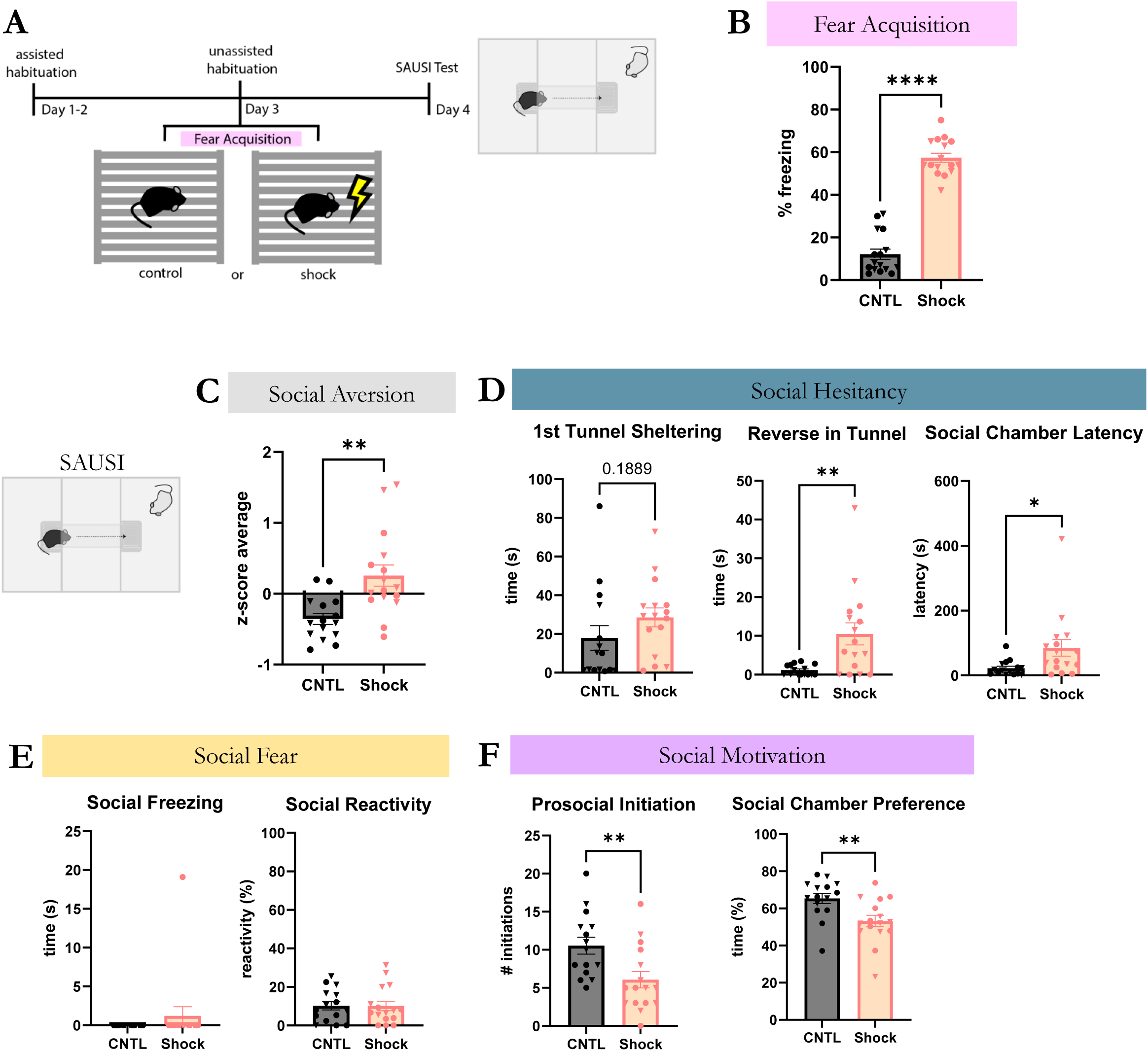
Foot shock stress-induced social aversion is detectable with SAUSI. (**A**) SAUSI timeline with foot shock stress. (**B**) During the foot shock protocol, mice that received the shock froze for significantly more time than controls. (**C**) In SAUSI, social aversion was significantly increased following foot shock compared to controls. (**D**) Social hesitancy behaviors – latency to enter the social chamber and reversing in the tunnel - were significantly increased by foot shock stress. (**E**) Foot shock stress did not impact social fear behaviors. **(F)** Social Motivation scores including time spent in the social chamber and number of prosocial initiations were decreased following foot shock. (n=8 females (circles), n=8 males (triangles) per group; independent samples t-tests, 2-tailed).

## DISCUSSION

Here we introduce SAUSI, a novel assay which can be used to assess social motivation, hesitancy, decision making, and in depth free social interaction in mice. SAUSI achieves this goal by enabling mice to choose between a “social” chamber and a “home” chamber using a tunnel for decisive access to either side, and combines this element of social choice and motivation with the ability to freely interact with a conspecific on the social side. Using this novel design, we wholistically characterize social aversion behavior, identifying critical features which drive this state: social fear, social motivation, and social hesitancy behaviors. SAUSI also enables us to assess generalized anxiety through non-social freezing and thigmotaxis behavior during baseline, and is compatible with USV recordings to monitor changes in vocal communication.

Our results demonstrate that isolation-induced changes in behavior are largely assay- and context-dependent. Certain behavioral assays or environmental contexts reveal territorial-driven changes to behavior following SI, while others are more sensitive to social motivation changes. Our results indicate that it is critical to choose an assay that accurately allows you to test your hypothesis, otherwise, competing behaviors or other environmental factors such as bedding may obscure findings. Existing assays such as the 3-chamber lack free behavior interaction, and a home cage testing removes the element of choice. Ultimately, this reveals SAUSI as a unique assay that is particularly fit to examine behaviors related to social aversion.

While we expected isolation to induce aggression in our RI assay^35,36,60,61^, we were excited to find SI-induced aggression in SAUSI as well. Two roles emerge in aggressive interactions - offensive and defensive. These have been described extensively^14,62–64^ and are indicated primarily by posture, advancement, and location of attack or submissive posture and flight behaviors, respectively. However, the circumstances and emotional states which motivate attack are less often discussed. Recent studies suggest that there may be a rewarding component for the aggressor which could provide motivation to initiate aggression (proactive rather than reactive aggression)^5,65–68^. Other theories suggest the establishment of dominance as key^14^.

SAUSI reveals that this isolation induced aggression is likely intrinsically motivated (proactive) in nature^5,65–68^. This is demonstrated by isolated mice repeatedly choosing to enter the social chamber in order to engage in immediate and repeated attack or mounting without provocation or reciprocation. Additionally, if aggression were reactive in nature, we would have expected to find similar rates of aggression towards a conspecific across all tests, especially when forced to interact with no escape options from the cage. However, we failed to see aggression in either the novel cage or no-bedding condition. We believe that the stark differences in aggression across contexts of forced interaction indicate that isolation-induced aggression is proactive, and likely driven by territorial attachment to a familiar environment. Our findings suggest that the motivation to be aggressive is intrinsically heightened for isolated mice, increasing the initiation of aggression altercations with a conspecific, unprovoked. On the other hand, foot shock stress did not cause an increase in aggression on SAUSI, which may lend to the possibility that foot-shock-induced aggression^69^ is more reactive in nature^5,65–68^. Further testing is needed to assess the role of dominance in influencing the motivation for aggression^14^. Additionally, studies that investigate aggression seeking behavior rely on learned reward tasks such as operant aggression seeking or aggression conditioned place preference, which require training and learning on behalf of the experimental mice^5,70^. SAUSI provides the ability to assess motivation for aggression in a more naturalistic setting, without training, and with free interaction.

In addition to nuances in motivation for aggression, SAUSI allows researchers to distinguish between different types of social aversion behaviors and across a wide range of subcategories. SI stress-induced social aversion is largely driven by social fear whereas FS stress-induced social aversion was more heavily influenced by reduced social motivation and changes in hesitancy and decision-making. This not only reveals social aversion as a whole, but can also provide insight into the underlying changes in emotional state that drive increased social aversion.

One intriguing discovery we made is that SAUSI allows for a number of behaviors to be displayed as a series along a continuum. Interestingly, this continuum seems largely governed by the imminence of the conspecific, consistent with theories which posit that animals will display a series of stereotypical and species-specific defense behaviors across a threat imminence continuum^71–74^. Indeed, when the distance between the experimental mouse and the conspecific mouse (or social “threat”) is large, animals display increased hesitancy behaviors. These can be thought of as examples of “pre-encounter” behavior, as mice are in a position to avoid potentially threatening social contact. As experimental mice gain proximity to the conspecific, but prior to contact, behavior continues to progress on a “social threat" imminence continuum. This can be indexed by increased hesitation to approach the social chamber, sheltering for long periods in the tunnel, and/or reversing out of the tunnel after the decision has been made to cross over to the social chamber. These behaviors may indicate an ongoing assessment of threat and incorporate retreat as the conspecific becomes closer. Finally, once mice enter the social chamber and begin to interact with the conspecific, encounter & circa-strike behaviors can be identified. These include social freezing and heightened social reactivity in response to being investigated. This continuum elucidated by SAUSI reveals the emergence of a state of social aversion^71–74^.

Ultimately, using new tools to investigate social aversion behavior is critical to understanding the neurobiology of associated disorders. Social anxiety, the fear of social situations, affects approximately 25 million adults in the United States alone^75,76^. It is estimated that over 5 million adults in the United States have been diagnosed with Autism Spectrum Disorder^77^. Each year autism and anxiety disorders have steadily risen^78,79^, particularly in the wake of the growing epidemic of loneliness^80^, suggesting that far more people are currently impacted. Despite the prevalence of social anxiety and autism spectrum disorder, our understanding of its etiology and underlying neurobiology is limited, hindered by the lack of animal models for crucial behavioral readouts, such as social aversion. Our findings impact the field of social anxiety and autism research by providing in-depth mouse models of stress induced social aversion behavior.

## MATERIALS AND METHODS

### Key resources table

| **Purpose** | **Resource** | **Link or Information** |
| --- | --- | --- |
| mice | C57BL/6J and BALB/c– Charles River Laboratories | https://emodels.criver.com/search?species=Mouse |
| cameras | Basler GigE camera acA1300-60gc | https://my.noldus.com/shared/knowledge-base/answer/256 |
| room setup | overhead pole for camera and lighting mount | Fudesy Photo Video Studeo 10Ft Heavy Duty Adjustable Backdrop Stand (Amazon) |
| room setup | blackout Curtains (Amazon) | Joydeco Blackout Curtains (Amazon) |
| room setup | anti-static spray | Plant Therapy Unscented Natural Wrinkle Releaser (Amazon) |
| lighting | gooseneck lamps with clamping base | GLFERA (Amazon) |
| microphone | avisoft | https://avisoft.com/ultrasound-microphones/cm16-cmpa/#40011 |
| SAUSI box and tunnels | CAD files (machine shop) | https://github.com/ZelikowskyLab/SAUSI_paper |
| foot shock board supplies | metallic tape | GENNEL 5mm x 20m Conductive Cloth Fabric Adhesive Tape (Amazon) |
| foot shock board supplies | buttons | Starelo 19mm 3/4" Momentary Push Button Switch (Amazon) – push & hold-ON; release-OFF |
| foot shock board supplies | alligator clip wires | Electronix Express 2 Wire 30 Ft Retractable Test Leads, 18 Gauge Wire with Alligator Clips (Amazon) |
| foot shock board supplies | power supply | Med Associates Fear Chamber and Aversive Stimulator/Scrambler https://med-associates.com/product/aversive-stimulator-scrambler-module/ |

### Data code and availability

The SAUSI apparatus is open-source, and design files are freely available online at: https://github.com/ZelikowskyLab/SAUSI_paper. In addition, we have made all analysis code for the figures in this paper available at https://github.com/ZelikowskyLab/SAUSI_paper.

### Subjects

One hundred C57Bl/6 mice and eighty BALB/c mice were housed in cages (Thoren type 9 https://thoren.com/?page_id=71) with Teklad pelleted bedding (https://www.inotivco.com/7084-pelleted-paper-contact-bedding) on a ventilated rack in a 12 hr light/dark cycle with full time access to food (Teklad Global Soy Protein-Free Extruded Rodent Diet 2920X) and water. All animal work was reviewed for compliance and approved by the Institutional Animal Care and Use Committee of The University of Utah (IACUC protocol #00001504).

### Design and construction of SAUSI

The SAUSI apparatus was designed in CAD and built by the machine shop at the University of Utah. The floors were made of white High Density PolyEthylene (HDPE) plastic, the tunnel (tube) and walls were made of transparent acrylic plastic. The tunnel block (5.3 x 2.5 x 6.3 cm) was made of metal (673 g - heavy material that mice cannot move), and the U-channel tunnel block was 3D printed. Transparent, plastic wall extenders (33 cm tall) were fitted to the exterior walls of the sausi box and secured with paper clips to ensure that mice could not jump out of the apparatus.

Foot shock boards were made by hand using metallic tape (optionally on a nonconductive backing (such as scotch tape) for portability). These were connected via alligator clip wires to the floor of the Med Associates fear chamber boxes. During the conspecific tunnel deterrent session, the footshock boards are wired directly to the fear chambers for automated shocking. During the SAUSI test day, this connection is mediated by off-on-off buttons for manual shock administration.

Black out curtains were hung surrounding the setup to prevent side preferences due to differences in visual stimuli across the room. They also improve the even distribution of lighting and increase sound dampening. Scent-free anti-static spray was sprayed on the footshock boards and curtains at the beginning of each day in the SAUSI protocol. This reduced the chances of accidental shock due to static.

### Room conditions

A bar was hung above the apparatus for mounting two yellow light lamps facing away from the mice and the single top view camera. Two side view cameras were also mounted to each side of the SAUSI box with permanent fixtures for consistency. The Avisoft microphone was clamped to the exterior middle chamber wall (Fig. 1a-b).

Lighting: 2 lamps facing upward, yellow hue, lux = 22-23 per chamber.

Temperature was 70-75 F in behavior suite, with air conditioning on (which provided low levels of background noise).

Handling with plastic beakers (4” diameter, 5” tall).

On the first day of the SAUSI protocol, mice were weighed. Each experimental mouse was paired with a lighter-weight BALB/c (2 weeks younger) that was sex matched. Each BALB/c was tested with both a GH and SI mouse in counterbalanced order. The hind region of BALB/c mice were colored with Stoelting animal markers (https://stoeltingco.com/Neuroscience/Animal-Markers~9769) to be identified throughout the training and testing.

### Mice

All mice were purchased from Charles River Laboratories. The strain of experimental mice were C57 BL/6, whereas the strain of conspecific were BALB/c mice. Experimental mice for adolescent isolation arrived at the University of Utah vivarium facilities at 5 to 6 weeks old (8 weeks old for foot shock experiments), with three to four mice per cage. All mice were housed on a ventilated rack with dark, opaque plastic dividers stationed in between cages. These features were present reduce the visual, auditory, and olfactory stimuli of surrounding cages.

### SAUSI protocol

The SAUSI apparatus contains two outer rooms, one empty and one containing a conspecific. These rooms are connected by a narrow tunnel with a miniature foot shock board on either side, which deter conspecifics from crossing over, and allows for observable decision making in the experimental mice.

To habituate mice to the apparatus, tunnel, and handling, experimental mice are placed in the arena for 5 minutes each on days 1 and 2. During this assisted habituation, experimenters gently guide the mouse through the tunnel every 30 seconds. Every 15 seconds, the experimenter places a hand in the opposite chamber of the mouse to habituate it to movement. This indicates to the mice that experimenters can move and intervene without attempting to handle the mouse every time and thus reduces intervention stress. Time restarts after each time the mouse crosses through the tunnel. For the very first entrance on the first day, mice are held by the tail with all 4 paws on the ground with their face directed near the entrance of the tunnel. From there, experimenters patiently wait until mouse enters the tunnel. Once all four paws are in, the mouse is blocked from reversing out of the tunnel ^37^. After the first entrance, “corralling” the mouse close to the tunnel entrance without touching it is sufficient for them to enter the tunnel and reduces stress. On the third day, mice are placed in the apparatus and monitored to ensure the mice use the tunnel at will (inspired by hierarchy tube habituation – fan et al). The conspecifics undergo a 5-minute foot shock deterrent session on day 3 to prevent them from crossing to the other side and blocking the tunnel entryway.

On day 4, the social behavior test consists of a 3-minute habituation period for the experimental mouse followed by a 10-minute test phase with the conspecific present (figure 1b). To begin the baseline phase, the tunnel is blocked off in the starting chamber. The experimental mouse is placed in one chamber (alternating between left and right for each trial) and the block is removed to signify the start of the 3 minutes. Once three minutes has past, the experimenter traps the mouse on the side it originally started on using the tunnel block. The conspecific is placed in the opposite chamber and the block is removed to signify the start of the 10-minute test phase. During the social test, the foot shock boards are controlled manually by a button and can be used if a conspecific steps on the foot shock board while the experimental mouse is not also on it. This only happens occasionally and can be reduced by minimizing the number of times each conspecific is used. To transport mice to and from the SAUSI apparatus, the tunnel is first blocked off and a plastic beaker (4” diameter, 5” tall) is used to transition the mouse back to its home cage.

For isolation, half of the mice were placed in social isolation (n=18 females, n=8 males), while the other half remained in their original group-housed cages (n=18 females, n=8 males). Isolation began at 6 weeks old and lasted for a total of 4 weeks (tested at 10 weeks old). Conspecifics were group-housed with 4 mice per cage and were tested at 8 weeks old (2 weeks younger than experimental mice). Each conspecific was paired with a GH and an SI mouse during the test day, counterbalanced for order.

### 3-Chamber

Lighting, video, and audio recording (as well as all other factors: curtains, wall extenders, anti-static spray, cup handling) were set up identically to SAUSI. The temperature was 70-75 F in the behavior suite, with air conditioning on (which provided low levels of background noise). The apparatus used for SAUSI was also used for 3-chamber testing, with the tunnel replaced by removable doors and with no foot shock boards present. Cups used to separate conspecifics (https://www.noldus.com/applications/sociability-cage) were placed in the top left or bottom right of the respective chamber and weighted down by water bottles. Videos were recorded from the overhead camera using MediaRecorder and scored using automated tracking in EthoVision. USVs were captured using Avisoft and were analyzed with DeepSqueak.

The 3-chamber task began with a 5-minute baseline phase, where experimental mice were placed in the middle chamber with nothing else present in the apparatus. The baseline phase began once the experimenter removed the doors, allowing the mouse to explore all 3 empty chambers. After the baseline phase, mice were then ushered back into the middle chamber, with access to other chambers blocked by doors in order to transition to the test phase. The test phase lasted 10 minutes and consisted of an inanimate object (black plastic block) under a cup in one chamber, and a conspecific mouse under a cup in the opposite chamber.

The “social” side was counterbalanced for each trial. Each conspecific was paired with a GH and an SI mouse during the test day, counterbalanced for order. The apparatus, cups, water bottles, and doors were cleaned with 70% ethanol between each trial.

For isolation, half of the mice were placed in social isolation (n=21 females), while the other half remained in their original group-housed cages (n=21 females). Isolation began at 6-14 weeks old and lasted for a total of 3-4 weeks. Conspecifics were group-housed with 4 mice per cage. Each conspecific was paired with a GH and an SI mouse during the test day, counterbalanced for order.

### Resident Intruder

Lighting was provided by two lamps facing upward with yellow hue, lux = 14-15. Temperature was 70-75 F in behavior the suite, with air conditioning on (which provided low levels of background noise). Videos were recorded from an overhead camera (43 cm from the cage floor) using MediaRecorder and manually scored in EthoVision. USVs were captured using Avisoft (microphone 38.1 cm from the cage floor) and were analyzed with DeepSqueak. A box surrounded each cage during testing and contained a ledge to prevent mice from exiting the cage during testing (https://github.com/ZelikowskyLab/SAUSI_paper).

The RI tests began with a 3-minute baseline phase, where the experimental mouse was observed in the cage on its own. This was followed by a 10-minute test phase, in which a conspecific mouse was placed in the cage with the experimental mouse.

Cages used (Thoren type 9- https://thoren.com/?page_id=71) were 19.56 x 30.91 x 13.34 (cm.) with a floor area of 435.7 (Sq. cm.). Bedding that mice were continuously housed in (Teklad pelleted bedding - https://www.inotivco.com/7084-pelleted-paper-contact-bedding) or new bedding of the same type were used for RI-home cage and RI-novel bedding. Nestlets were removed for visibility throughout the task. Food and water were not accessible during the baseline and test phases of the task.

Group-housed mice that were not actively being tested were placed together in a clean holding cage. Each group-housed mouse shared an “untested” holding cage with their cage mates prior to testing. After being tested, they were placed in another clean “tested” holding cage so that no untested mice were exposed to tested mice.

For isolation, half of the mice were placed in social isolation (n=8 females), while the other half remained in their original group-housed cages (n=8 females). Isolation began at 6 weeks old and lasted for a total of 4 weeks (tested at 10 weeks old). Conspecifics were group-housed with 4 mice per cage and were tested at 8 weeks old (2 weeks younger than experimental mice). Each conspecific was paired with a GH and an SI mouse during the test day, counterbalanced for order.

### Foot Shock Stress Paradigm

All mice (shock and controls) were isolated for 7 days prior to the SAUSI test day to prevent within-cage aggression following shock, and to not reach the threshold for chronic social isolation (2+ weeks)^61^. On the third day of the SAUSI timeline, after receiving the unassisted tunnel habituation, each mouse was placed in a Med Associates fear chamber. Mice in the shock group (males, n=8; females, n=8) received 10 1-second, 1-mA pseudorandomized foot shocks over a 60 minute period^59^. Controls (males, n=8; females, n=8) were placed in the Med Associates fear chambers as well, but received no shocks over the 60 minute duration.

Conspecifics were group-housed with 4 mice per cage and were tested at 8 weeks old. Each conspecific was paired with a CNTL and an Shock mouse during the test day, counterbalanced for order.

### Behavior and Vocalization Analysis

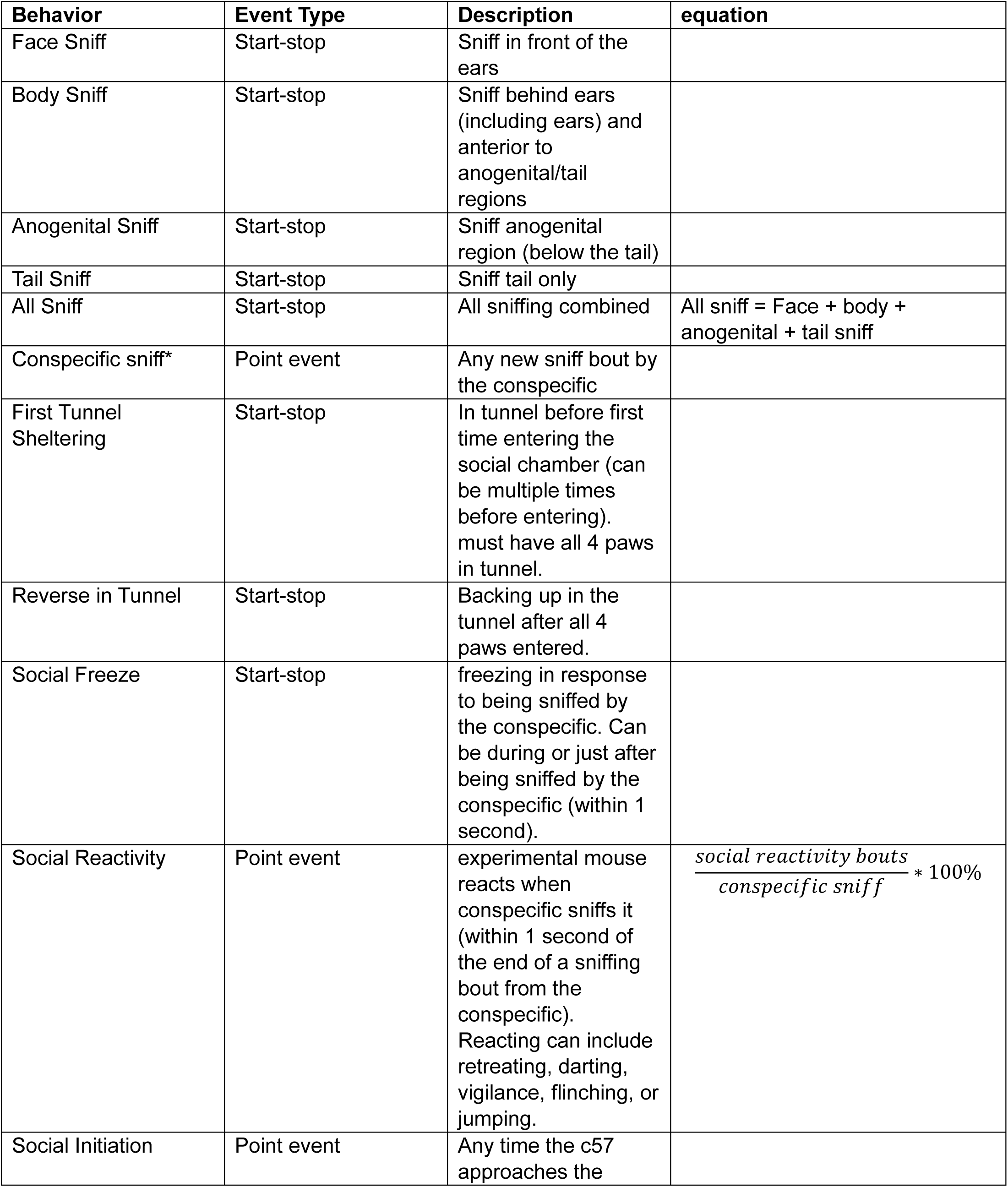
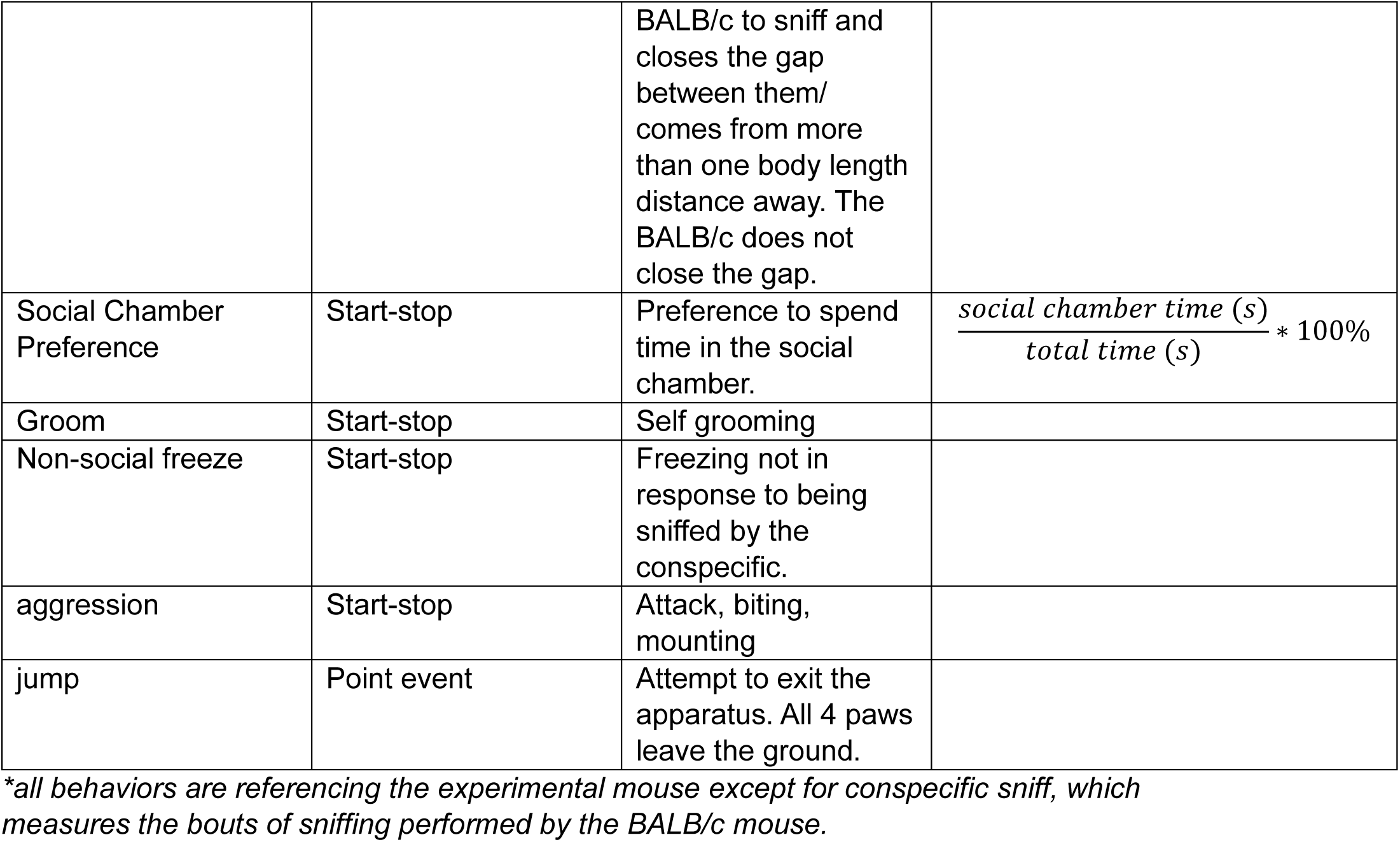

Videos were recorded in MediaRecorder. The top view perspective was used for automated analysis latency and time spent in each chamber with EthoVision^81^ as well as postural data tracking in SLEAP^17^. Side view perspectives were used for hand scoring behaviors in EthoVision.

DeepSqueak was used to analyze all audio recording data from avisoft.

SLEAP ^17^ was the program used for postural tracking. All videos used for tracking were from the top camera view and were trimmed to only include the test phase. These videos were used for training in SLEAP with >600 frames manually labelled. The “export labels package” feature from the SLEAP user interface was used to export labelled frames, which were uploaded in google colab, where training was performed (Google Colab file accessible using the “train on google colab” feature in the SLEAP user interface). Bottom-Up processing was used for training with default settings. For inference, tracker was set to “flow”, clean_instance_count was set to 2, track_window was set to 30, and all remaining settings were left as default. Google Colab was used for training and inference due to high computational needs.

The Motion Mapper pipeline^21^ was used for unsupervised behavioral analysis. (modified code available here https://github.com/ZelikowskyLab/SAUSI_paper).

Python and matlab were both used to generate nose to nose distance calculations, behavior rastor plots, all and all regression data (code available here https://github.com/ZelikowskyLab/SAUSI_paper).

GraphPad PRISM was used to generate all statistics and graphs comparing GH and SI conditions.

### Exclusion Criteria

1 mouse was excluded from the figure 1d data because the conspecific stepped on the footshock board (and received a shock) during the first tunnel entrance of the experimental mouse, which can impact latency to approach, first tunnel sheltering, and reversing in tunnel. Other pre-defined exclusion criteria for SAUSI include the experimental mouse failing to use the tunnel by the 3^rd^ habituation day, receiving an accidental foot shock, or conspecific aggression. None of these criteria were met throughout experimentation.

Motion map regions 1 and 2 were excluded from logistic regression analyses (Fig. 2e) because they represent outlier/missing tracked points from SLEAP.

3 mice were excluded from the social interaction preference and social approach preference data in the 3-chamber experiment (Fig. 3a) because they climbed on top of the 3-chamber cups during the test phase and an experimenter intervened.

Predefined exclusion criteria for RI experiments included BALB/c aggression. 4 mice were excluded from figure 3 “# USVs” data due to BALB/c aggression. All other scores include data from these mice only prior to the first bout of BALB/c aggression.

1 mouse was excluded from the figure 4 data due to injury just prior SAUSI testing.

## ACKNOWLEDGEMENTS

We thank the University of Utah Machine Shop for building the SAUSI apparatus; Adilenne Maese for running and scoring the home cage resident intruder assay; Dr. Sewon Park for providing Matlab code for nose-to-nose distance analysis; Alan Mo for GitHub management; Dr. Neda Nategh for advice about model fitting; and Benjamin Dykstra and Kanishk Jain for trouble shooting motion mapper. This work was supported by the National Science Foundation Graduate Research Fellowship Program (JG), Travel scholarship to The Short Course on the Application of Machine learning for Automated Quantification of Behavior, Jackson Laboratory, ME (JG), an NIMH R01 MH132822 (MZ), a Klingenstein-Simons Early Investigator Award (MZ), a Whitehall Fellowship (MZ), a Sloan Fellowship (MZ), a McKnight Scholars Award (MZ), and the University of Utah.

## Notes

https://github.com/ZelikowskyLab/SAUSI_paper

### Competing Interest Statement

The authors have declared no competing interest.

### Summary of Updates

updated all main and supplemental figures, included additional foot shock stress data, and broadened the scope to include social aversion more generally

