## supplemental figures for "SAUSI: an integrative assay for measuring social aversion and motivation"

### Figure 1 – figure supplement 1

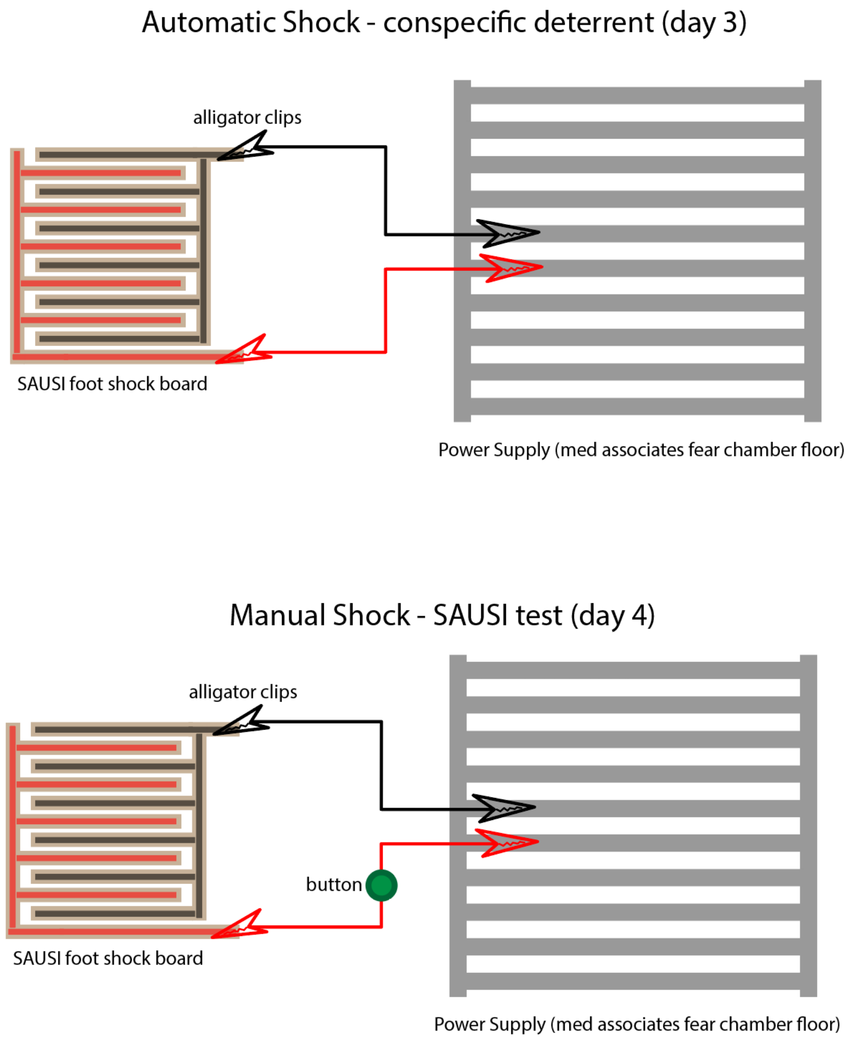

### Figure 1 – figure supplement 2

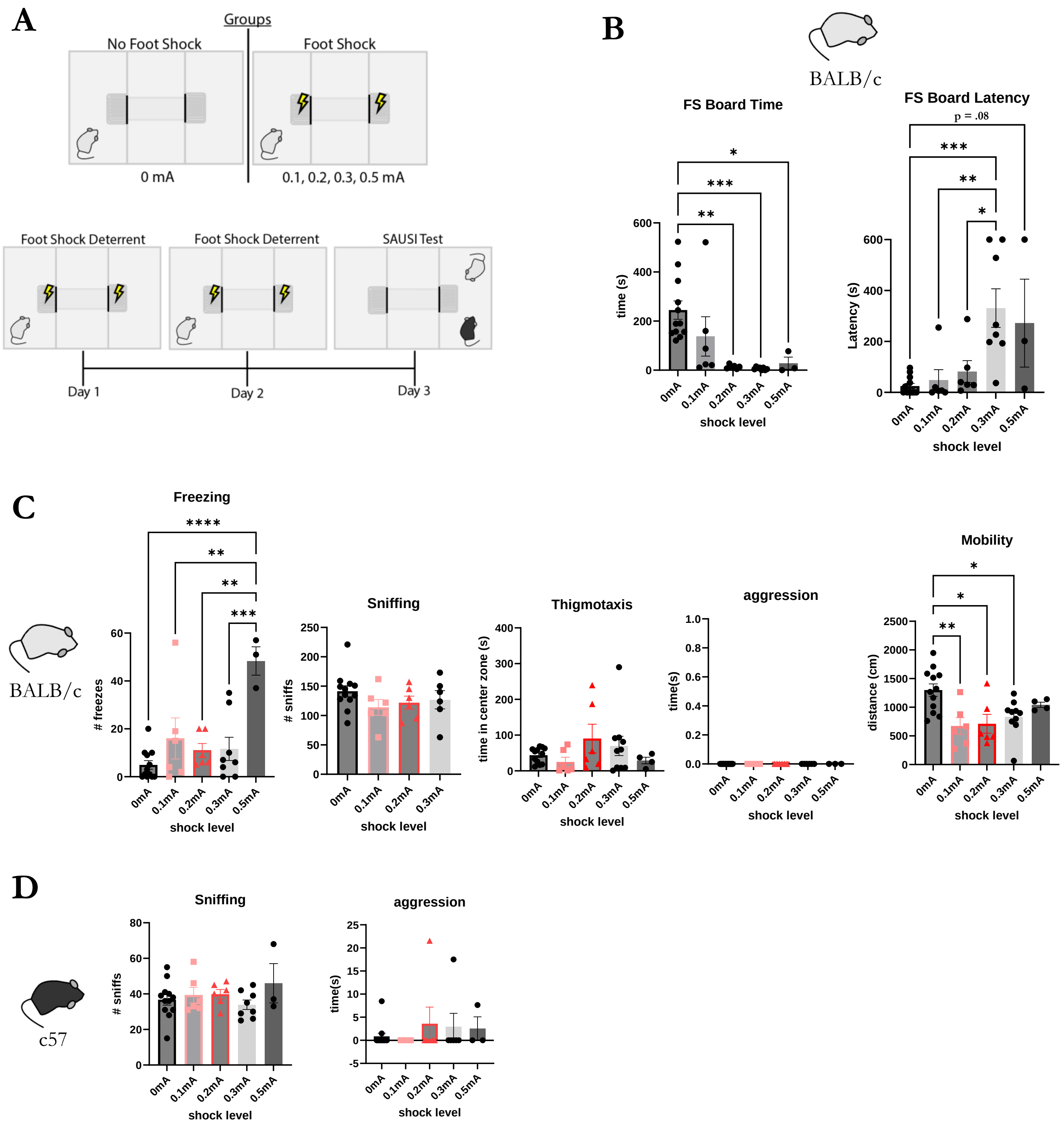

Figure 1 – figure supplement 3

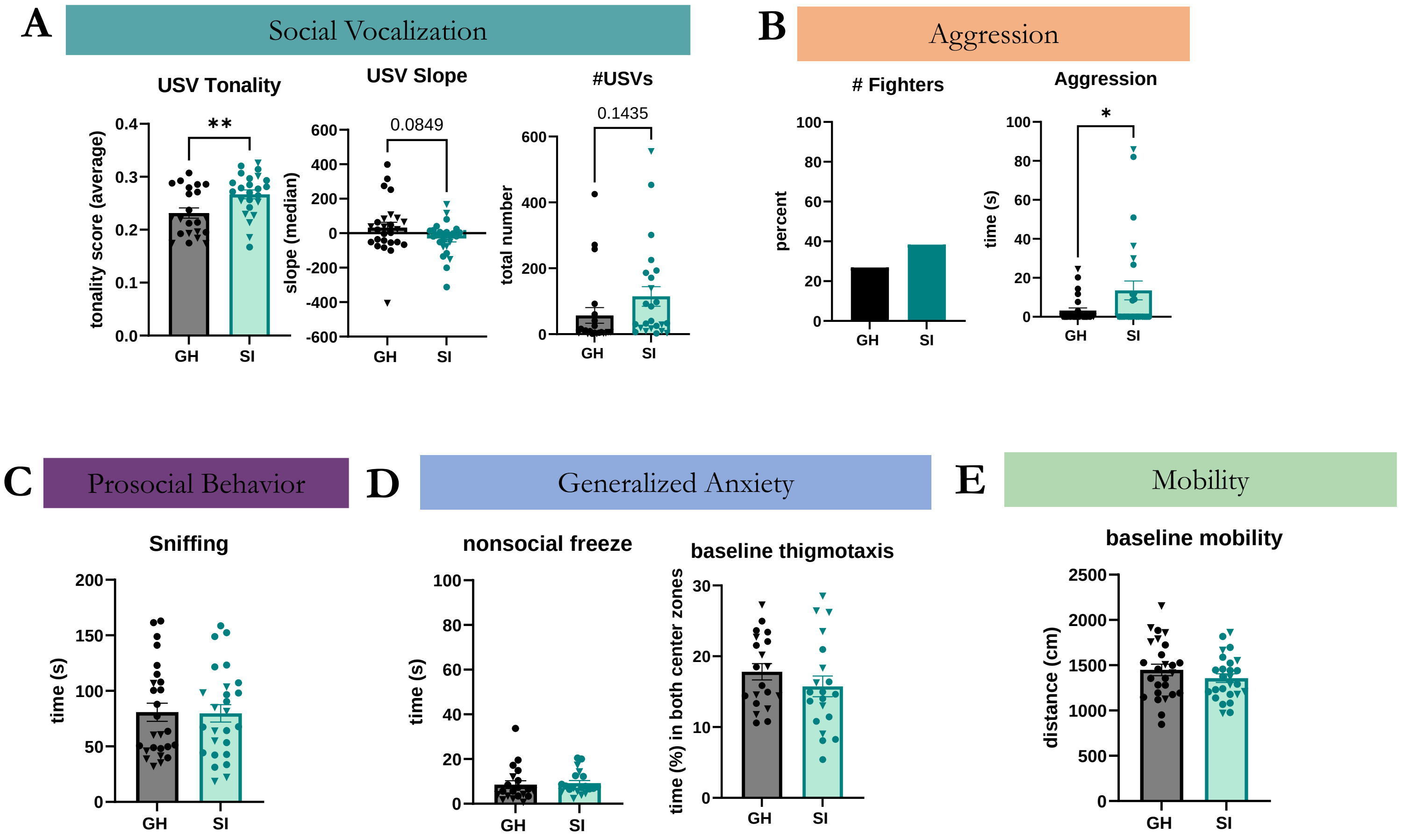

### Figure 3 – figure supplement 1

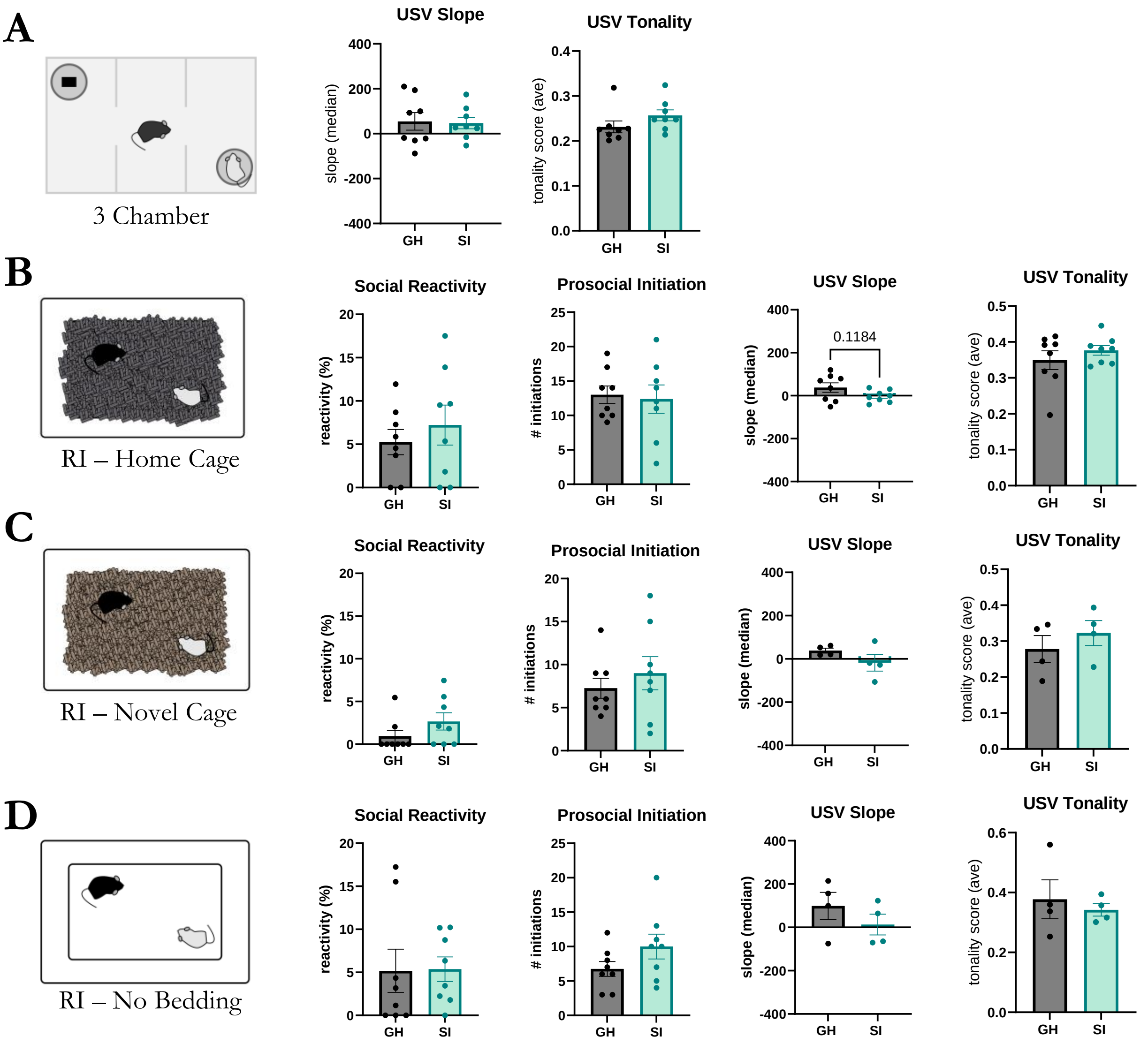

Figure 4 – figure supplement 1

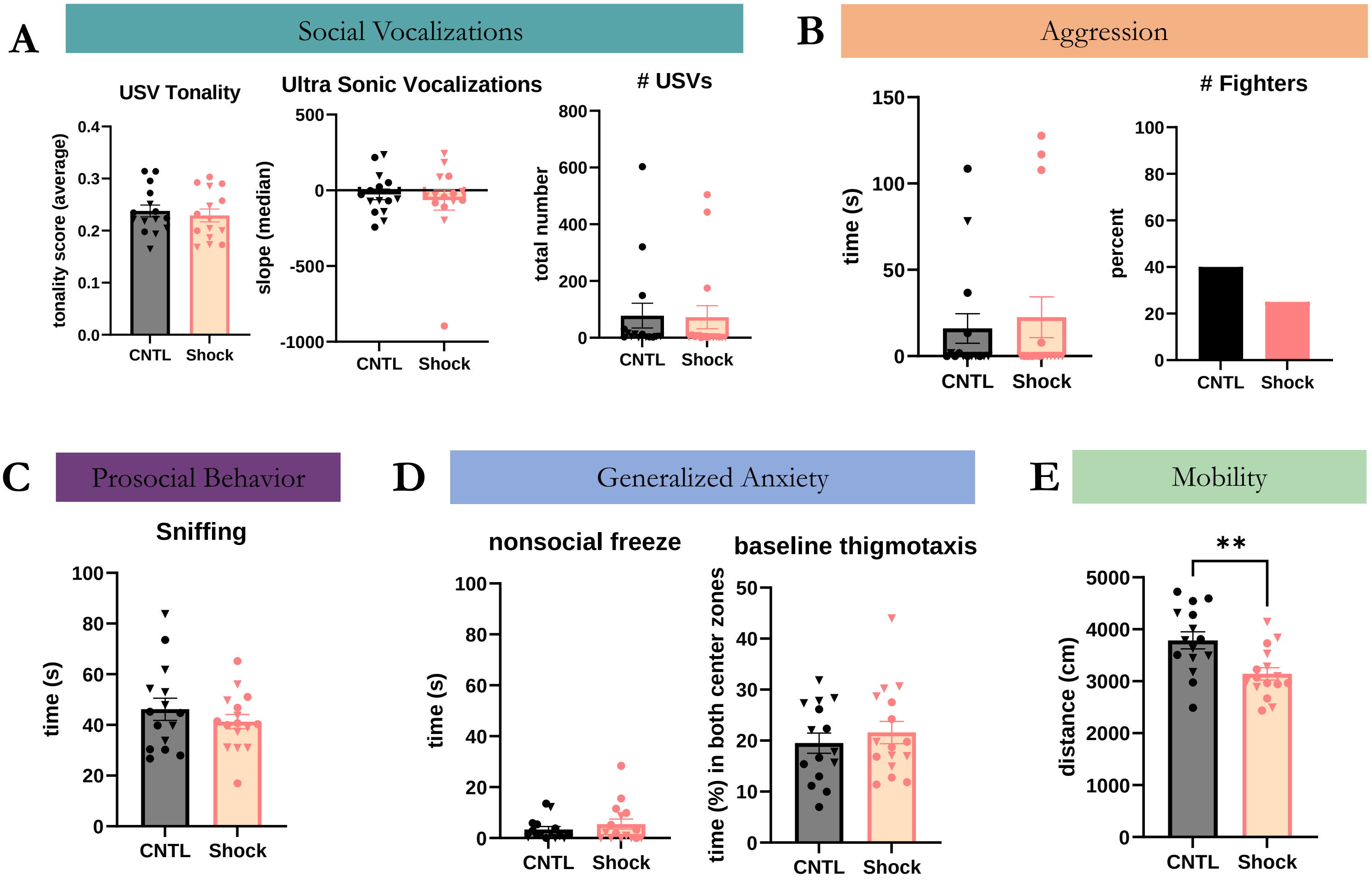
